# Sublethal effects of the organosilicone surfactant Silwet L-77 and its interactions with acetamiprid and sulfoxaflor on honeybee (*Apis mellifera*) gustatory responsiveness and olfactory learning

**DOI:** 10.64898/2026.06.28.735033

**Authors:** Maja Achatz, Nadine Benda, Magdalena M. Mair, Julia Osterman, Christoph Kurze

**Affiliations:** Evolutionary Ecology, University of Regensburg, Germany; Statistical Ecotoxicology, University of Bayreuth, Germany; Department of Biological and Environmental Sciences, University of Gothenburg, Sweden; Department of Biological and Environmental Sciences, Gothenburg Global Biodiversity Centre, Sweden

**Keywords:** olfactory conditioning, gustatory responsiveness, honey bee, Silwet, neonicotinoid, sulfoximine

## Abstract

Pesticides remain indispensable for global food security, yet their use must be reconciled with the preservation of biodiversity. Despite advances in developing ‘safer’ pesticides, their sublethal effects and synergistic potential with co-formulants in tank-mixtures, including ‘inert’ spray adjuvants, often remain poorly understood in beneficial insects like bees. In this study, we assessed the effects of acute oral exposure to both single compounds and adjuvant-insecticide mixtures on gustatory responsiveness, associative olfactory learning, and 90-minute short-term memory retention in honeybee foragers (*Apis mellifera* L.). Specifically, individual foragers were exposed to either single compounds (0.1%, 0.5%, 1% or 2.5% Silwet^TM^ L-77; 6, 30 or 150 ng acetamiprid; 3, 9 or 27 ng sulfoxaflor), adjuvant-insecticide mixtures (0.5% or 1% Silwet^TM^ L-77 combined with 6 or 30 ng acetamiprid, or 3 or 9 ng sulfoxaflor), a positive control (6 ng imidacloprid), or negative controls (0%, 0.1%, and 1% acetone). While we found no significant interaction effects for these adjuvant-insecticide mixtures, single-compound exposure to the highest doses of 2.5% Silwet^TM^ L-77 and 27 ng sulfoxaflor significantly reduced gustatory responsiveness. Furthermore, all three agrochemicals alone weakly affected associative learning without impacting short-term memory retention. Although adjuvant-insecticide mixtures did not significantly impair learning ability, these adjuvants may impact sensory perception. Moving beyond mortality-based assessments and including behavioural assay to test sensory perception and learning ability is essential for gaining a more holistic understanding of pollinator health.

**Graphical abstract:** 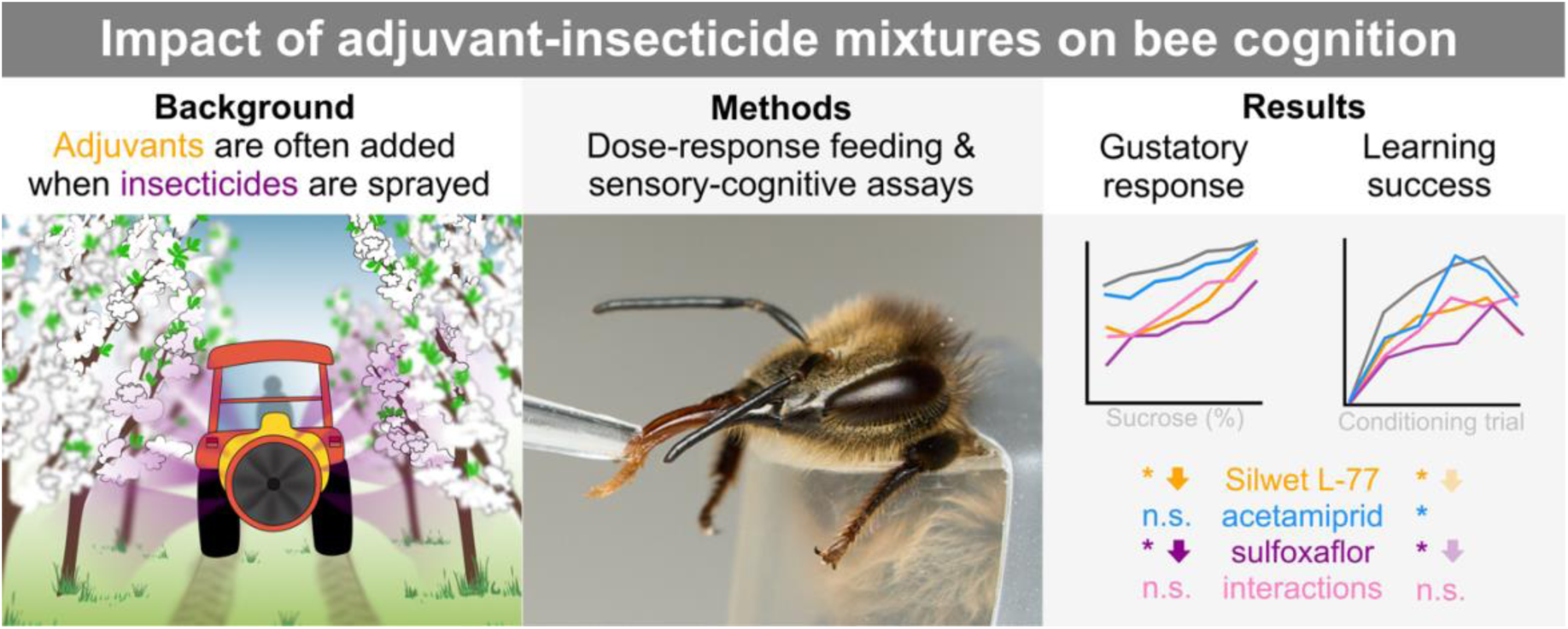

## 1. Introduction

The widespread application of agrochemicals has been identified as a key factor contributing to the concerning decline of insect pollinators, alongside habitat loss, climate change, and the spread of emerging pathogens (e.g. Dicks et al., 2021; Goulson et al., 2015; Potts et al., 2010). Until the last decade, neonicotinoids were the most widely used insecticides worldwide, but their impact on non-target pollinators, including bees, was alarming and called for rapid regulatory action (Dentzman et al., 2025; Simon-Delso et al., 2015). By targeting nicotinic acetylcholine receptors (nAChRs) in the central nervous system in insects, these systemic compounds also caused sublethal effects in bees such as impaired cognition, altered foraging behaviour, and decreased fecundity (e.g. Mandal et al., 2026; Spence et al., 2026). To mitigate these risks, the European Union (EU) banned legacy neonicotinoids such as imidacloprid (EC, 2018a, b, c) to initiate an agricultural shift towards ‘bee-safe’ alternatives such as acetamiprid (ACE; a cyano-substituted neonicotinoid) and sulfoxaflor (SFX; a sulfoximine). These compounds are characterised by a lower acute toxicity to pollinators, specifically honeybees, compared to legacy neonicotinoids (e.g., DesJardins et al., 2023; Mandal et al., 2026). While these regulations led to a notable decrease in the use of legacy insecticides, the risk of unregulated chemical substitutes and lesser-understood alternatives remains (Gensch et al., 2024). For example, SFX was subsequently banned for outdoor use in the EU (EC, 2022).

A major limitation of current risk assessments is the reliance on mortality assays rather than integrating the assessment of sublethal effects that are critical indicators for the long-term sustainability (Desneux et al., 2007; Tosi et al., 2022). To make matters worse, active ingredients are rarely applied in isolation. Instead, they are used in combination with co-formulants in tank-mixtures, including adjuvants such as Silwet^TM^ L-77 (SIL, also known as “the super spreader”), which are applied to enhance pesticide penetration and efficacy (Mullin et al., 2016; Shannon et al., 2023; Straw et al., 2022). Despite being labelled as ‘inert’, these adjuvants reduce surface tension by design and may bypass protective cuticular barriers and the insect intestinal epithelium, potentially inducing unexpected synergistic toxicities with ‘safe’ pesticides (Shannon et al., 2023; Straw et al., 2022).

Classical conditioning using the proboscis extension response (PER) assay has become the gold standard in ecotoxicology for assessing sublethal effects on learning and memory retention in honeybees, *Apis mellifera* L. (Hymenoptera: Apidae), due to its high sensitivity and reliability (DesJardins et al., 2023; Matsumoto et al., 2012; Scheiner et al., 2013). While extensive research demonstrates that legacy neonicotinoids (e.g., imidacloprid, thiacloprid, and clothianidin) significantly impair cognition, the effects of acetamiprid are inconsistent (e.g., DesJardins et al., 2023; Mandal et al., 2026; Simon-Delso et al., 2015). For example, although evidence suggests that acetamiprid has no significant impact on PER learning in honeybee foragers either as a pure substance (Aliouane et al., 2009; El Hassani et al., 2008) or as a commercial formulation (Schuhmann and Scheiner, 2023), decreased learning ability has been observed in young workers (Shi et al., 2019). Furthermore, acetamiprid has been shown to increase sucrose sensitivity (gustatory response), impair memory capacity, decrease homing ability, and alter foraging behaviour in honeybee workers (El Hassani et al., 2008; Shi et al., 2019; Shi et al., 2020), though these findings were not confirmed in other studies (Aliouane et al., 2009; Capela et al., 2022). Studies concerning sublethal doses of sulfoxaflor also indicate impaired learning (Cartereau et al., 2022) and homing ability (Capela et al., 2022) in honeybees, while other studies found no impact on olfactory learning in honeybees or bumblebees (Siviter et al., 2019; Vaughan et al., 2022). While acetamiprid and sulfoxaflor may be less harmful individually, there is a substantial risk that they synergistically interact with other agrochemicals such as spray adjuvants (e.g., Chen et al., 2019; Straw et al., 2022; Wernecke et al., 2022). Despite this concern, to our knowledge, only one study has investigated the effects of commonly used spray adjuvants on bee olfactory learning. Specifically, Ciarlo et al. (2012) showed that 1% Silwet^TM^ L-77, Dyne-Amic^®^, Syl-Tac^®^, and Sylgard^TM^ in sucrose solution impaired PER learning and short-term memory retention in honeybees.

To bridge the knowledge gap regarding the acute oral exposure to sulfoxaflor, acetamiprid, spray adjuvants, and their mixtures at potential field concentrations, further attention is needed to inform policy makers. While PER learning and memory retention assays standardise the assessment of the sublethal effects of agrochemicals on honeybee cognition, the gustatory response score (GRS) is a crucial additional method that is often ignored. In fact, sucrose responsiveness provides fundamental insights into learning and foraging behaviour (Scheiner et al., 2013). For example, if a pesticide, surfactant, or their interaction masks the perception of food, sublethal effects on learning may manifest as a downstream result of altered foraging activity. In this study, we address this knowledge gap by evaluating the effect of sublethal (field-relevant) doses of acetamiprid, sulfoxaflor, and their interactions with the common spray adjuvant Silwet^TM^ L-77 on sensory perception and learning ability. Using a dose-response framework incorporating GRS and PER learning assays, we aimed to answer the following research questions: (1) What are the acute effects of single-compound oral exposure to Silwet^TM^ L-77 (0.1%, 0.5%, 1%, 2.5%), acetamiprid (6 ng, 30 ng, 150 ng), and sulfoxaflor (3 ng, 9 ng, 27 ng) on sucrose responsiveness and learning behaviour? (2) Are there acute interaction effects of adjuvant-insecticide mixtures at potential field-relevant doses (0.5% or 1% Silwet^TM^ L-77 combined with 6 ng or 30 ng acetamiprid, or 3 ng or 9 ng sulfoxaflor)?

## 2. Materials and methods

### 2.1 Experimental overview

To test the acute sublethal effects of Silwet L-77 (SIL) and its synergistic potential with acetamiprid (ACE) and sulfoxaflor (SFX) on honeybees (*Apis mellifera* L.), we used a total of 862 foragers sourced from four queenright colonies. An overview of the daily experimental routine is outlined in the supplementary material (Fig. S1a). Briefly, collected bees were randomly allocated into two sequential groups, with the second group undergoing the identical experimental procedure as the first, but with a 70-min delay for practical reasons. First, bees were harnessed and allowed to habituate for 120 min in an incubator (Binder KT 115, Binder GmbH) at 25 ± 1°C with 50-60% RH to induce a standardised baseline state of starvation. Prior to receiving one of the randomly and blindly assigned treatments (Table 1), each bee was pre-tested for hexanal (neutral odour stimulus) and 30% sucrose to assess its baseline spontaneous response and motivation, respectively. Following oral treatment, bees were returned to the incubator for a 90-min exposure period to allow compound absorption prior to testing. Subsequently, we conducted the Gustatory Response Score (GRS) assay and Proboscis Extension Response (PER) olfactory conditioning assay following standard procedures (Scheiner et al., 2013). Short-term memory retention was assessed after a final 90-min resting period.

**Table 1.** Overview of experimental design. Due to the complexity of the dose-responses and interactions, experiments were conducted in three experimental blocks (EB). Each bee was fed its respective treatment dose in 2 μL of 50% sucrose solution at the start of the experiment. Sample sizes (N) indicate the total number of bees per treatment at the end of the GRS (Gustatory Response Score) and PER learning (Proboscis Extension Response) assays. The total number of bees that died (DIED) or showed any abnormalities (ABN) per treatment during the experiments. Differences in N between the GRS and PER learning assay reflect either bees that died or bees unresponsive to the 30% sucrose stimulus either due to the lack of motivation or sensory perception. Hence, those were removed from the statistical analysis of the PER learning data. In rare cases bees escaped (prior GRS: 1 bee of SIL 0.1%, 1 of SIL 0.5%, 1 of SIL 1%, and 2 of SIL 2.5% treatment).

| Treatment | EB<sup>‡</sup> | Doses | N (GRS) | N (PER<sub>L</sub>) | DIED | ABN<sup>§</sup> |
| --- | --- | --- | --- | --- | --- | --- |
| Ctrl (control) <sup>#</sup> | 1 | 50% w/w sucrose | 48 | 48 | 1 | 0 |
| Acetone Ctrl <sup>#</sup> | 1-3 | 0.1%, 1% v/v | 32, 82 | 32, 81<sup>“</sup> | 0, 0 | 0, 0 |
| IMI (Imidacloprid) | 1 | 6 ng | 50 | 38 | 0 | 3<sup>°</sup> |
| SIL (Silwet<sup>™</sup> L-77) | 1 | 0.1%, 0.5%, 1%, 2.5% | 48, 49, 49, 48 | 48, 48, 48, 45 | 1, 0, 1, 0 | 0, 0, 0, 0 |
| ACE (Acetamiprid) | 3 | 6 ng, 30 ng, 150 ng | 32, 32, 31 | 32, 32, 30 | 0, 0, 2 | 0, 0, 0 |
| ACE + SIL 0.5%<sup>*</sup> | 3 | 6 ng, 30 ng | 32, 32 | 32, 32 | 0, 0 | 0, 0 |
| ACE + SIL 1% | 3 | 6 ng, 30 ng | 32, 31 | 32, 31 | 0, 1 | 0, 0 |
| SFX (Sulfoxaflor) | 2 | 3 ng, 9 ng, 27 ng | 32, 32, 31 | 32, 32, 22 | 0, 0, 8 | 0, 0, 2<sup>§</sup> |
| SFX + SIL 0.5%<sup>*</sup> | 2 | 3 ng, 9 ng | 32, 32 | 30, 32 | 0, 0 | 1<sup>§</sup>, 0 |
| SFX+ SIL 1% | 2 | 3 ng, 9 ng | 32, 32 | 32, 31 | 0, 1 | 0, 0 |

<sup>#</sup> Controls were pooled as they differed not significantly from each other (see results; Figs. 1a, 2a)
<sup>\*</sup> Results are presented in supplementary Figs. S2 and S3
<sup>‡</sup> Experiments were conducted in June/July (Block 1), July (Block 2), and August (Block 3)
<sup>“</sup> One bee escaped during experiment
<sup>§</sup> Number of bees showing abnormalities per treatment (<sup>°</sup> lethargic; <sup>§</sup> twitching or moving a lot)

### 2.2 Preparation of treatment solutions

Twenty-two treatments were freshly prepared in 50% w/w sucrose solution prior to the experiment and stored as aliquots at -20°C (Table 1). Control bees (Ctrl) received either a pure 50% sucrose solution, or 50% sucrose solution supplemented with 0.1% or 1% acetone (ACS reagent, ≥ 99.5% (GC), CAS: 67-64-1, Sigma-Aldrich Chemie GmbH) to account for potential solvent effects, as all insecticides required initial dissolving in 1 mL acetone before further dilution. While 6 ng (i.e. 3 ng μL^-1^) imidacloprid per bee (IMI; Pestanal^®^, CAS: 138261-41-3, Sigma-Aldrich Chemie GmbH) served as a positive control, concentrations of acetamiprid (ACE; 6, 30, and 150 ng/bee, Pestanal^®^, CAS: 160430-64-8, Sigma-Aldrich Chemie GmbH) and sulfoxaflor (SFX; 3, 9, and 27 ng/bee, Dr. Ehrenstorfer^TM^, CAS: 946578-00-3) being the insecticides in focus of this study. Silwet L-77 (SIL; Silwet^TM^ L-77 Super Spreader, CAS: 27306-78-1, provided by Momentive Performance Materials GmbH) was diluted to 0.1%, 0.5%, 1%, and 2.5% v/v in 50% sucrose solution. Finally, treatment combinations were prepared containing either ACE (6, 30 ng/bee) or SFX (3, 9 ng/bee) both with 0.5% and 1% SIL.

### 2.3 Collection of test bees

We collected foragers from four honeybee hives (*Apis mellifera*), kept at the Botanical Garden at the University of Regensburg (Germany), from 26^th^ of June to 16^th^ of August 2024. Sampling rotated between hives daily to ensure balanced sample sizes across colonies, treatments and environmental variation. Both returning and leaving foragers were carefully collected with soft tweezers at the hive entrance in the morning between 8:30 and 9:00 and randomly split into two sequential groups, each separately kept inside a stainless-steel cage (12.4 x 5.8 x 15 cm) with access to 30% w/w sucrose solution *ad libitum*. Subsequently, honeybees were transported to the laboratory, where we conducted all experiments under standardised laboratory conditions.

### 2.4 Harnessing

Twenty honeybees of the first group were individually transferred to 20 mL glass vials and briefly immobilized on ice for about 2-3 min, while the second cage remained in the incubator. Each bee was harnessed in a 1.5 mL centrifuge tube with the bottom cut off, allowing the bee’s head and two front legs to pass through the hole. Bees were secured with a strip of duct tape (see Fig. S1b). Their head and two front legs remained unrestricted in movement. A small piece of PU foam (made from hybrid plugs, *Drosophila* Center, K-TK e.K.) was placed inside the tube to prevent bees from slipping out. Harnessed bees were starved for 2 hours in the incubator to ensure successful treatment and a high motivation during the entire assays. The second group underwent the same procedure as the first group, but with a 70-minute delay for practical reasons.

### 2.5 Pre-tests

We tested each bee’s responsiveness to the neutral stimulus to reduce physiological and behavioural variation among individuals following standard procedures (Scheiner et al., 2013). Specifically, each bee was exposed to the neutral odour stimulus hexanal (98% GC, CAS: 26-25-1, Sigma-Aldrich Chemie GmbH) for 6 s in the same experimental setup as described below to assess the unconditioned PER baseline response and serving as a negative control. Secondly, each bee was presented a toothpick dipped in a 30% w/w sucrose solution near their antennae to elicit a PER serving as a pre-treatment positive control and indicator of the bee’s physiological state and motivation (Scheiner et al., 2013). Following standard procedures, any bee responding the neutral odour (< 5%) or being unresponsive to 30% sucrose solution (< 20%) were excluded from the experiment (Bitterman et al., 1983; Tan et al., 2015). From the remaining bees, we randomly selected 14-16 bees per sequential group for the subsequent treatment and behavioural assays.

### 2.5 Treatment

Honeybees were randomly assigned to blinded treatments, accounting for any potential experimental biases, and fed a 2 μL droplet sucrose solution containing the respective dose (Table 1, Fig. S1b). In the extremely rare case that a bee did not consume the entire droplet, this individual was excluded from experiment. As potential daily field exposure levels have been reported to range from 6.4 to 115 ng/bee/day for ACE and from 5.8 to 30.06 ng/bee/day for SFX (Capela et al., 2022), we chose 6, 30, and 150 ng ACE and 3, 9, and 27 ng SFX as oral exposure doses, with the highest doses representing the extreme exposure scenarios. Although manufacturer recommendations for SIL in tank-mixes often range between 0.015% and 0.15%, we utilized a range of 0.1% to 2.5% v/v for three reasons: (1) manufacturer recommendation in ready-to-use agrochemical formulations is ≥ 5%, (2) bees in this study received only a single acute dose in 2 µL whereas bees in natural environments may feed repeatedly on treated crops, and (3) environmental concentrations on floral surfaces can increase significantly following a spray event due to the rapid evaporation of water (Straw et al., 2022). Following the feeding, honeybees were returned to the incubator for 90-minutes to allow compound absorption prior testing similar to previous research (Tan et al., 2015) and for practical reasons. Due to the complexity of the dose-responses and interactions, experiments were conducted in three experimental blocks, each lasting for three weeks. Block 1 served to establish the baseline for SIL dose-responses and positive controls. Block 2 and 3 focused on the acetamiprid and sulfoxaflor, respectively, including their dose-responses and interactions with SIL.

### 2.5 Experimental setup

Our setup (detailed in Fig.S1b) consisted of a polystyrene box (21.5 x 21.5 x 21.5 cm) used as the conditioning arena, which was closed with an acrylic glass lid. A continuous stream of air (1.2 L min^-1^; stimulus controller CS-55, Ockenfels Syntech GmbH) was delivered through one of two disposable Pasteur pipettes (air only), both passing through the wall of the polystyrene box. For the pre-conditioning test and for PER learning trials, the air stream was redirected through the second glass pipette for 6s, which contained 4 µL hexanal on a 4 x 1 mm piece of filter paper (neutral odour stimulus). Each test bee was placed approximately 4.5 cm in front of the of these pipettes (Fig. S1d). A table fume hood (Siemtech Elektrotechnik Heuschmann GmbH) was positioned behind the bee to exhaust the air stream. The sucrose reward was provided through a small window (5.5 x 7 cm) at the front of the polystyrene box.

### 2.5 Gustatory responsiveness assays

Prior to classical olfactory conditioning, the gustatory responsiveness of each bee was tested following standard protocols (Scheiner et al., 2013). Briefly, increasing concentrations of sucrose solution (0% (tap water), 0.1%, 0.3%, 1%, 3%, 10%, and 30% w/w) were individually presented to their antennae with a toothpick that were soaked in the respective sucrose solution at 2-minute intervals. The PER (Proboscis Extension Response) for each bee and concentration was recorded to calculate the Gustatory Response Score (GRS), with a positive response (spontaneous, unconditioned PER) scored as 1 and no extension as 0. These binary scores were subsequently used to determine the GRS proportions per treatment group. Any bee unresponsive to 30% sucrose was excluded from further testing to ensure only motivated and sensory perceptive individuals were included.

### 2.6 Learning and short-term memory assays

Learning ability and 90-minute short-term memory retention were assessed following standard procedures for classical olfactory conditioning (Scheiner et al., 2013). Briefly, individual bees were conditioned to a neutral odour (hexanal) over five acquisition trials (PER learning/conditioning) at 10-minute intervals. For this, each harnessed bee was put into the conditioning arena. After an initial 10-second acclimatization period (air stream at 1.2 L min^-1^), each bee was then exposed to an air stream with hexanal for 6 s per conditioning trial (Fig. S1c). During the first half (3 s), no reward was provided (testing phase; see positive PER in Fig. S1d). During the second half of hexanal exposure (lasting also 3 s), each bee was rewarded (conditioning phase) by allowing them to briefly antennate and feed from a toothpick soaked in 30% w/w sucrose solution. After every conditioning trial, the individual was removed, the conditioning arena was vented for 18 s, and the next bee was placed inside. To minimize experimental bias, the treatments were blinded and their test order was randomized daily. The sequential testing order of individual bees was maintained across all conditioning trials to ensure consistent intervals of 10 min for each bee. After each conditioning interval, the filter paper with 2 µL hexanal was renewed.

After completing the fifth trial, all bees were fed 5 µL of 30% sucrose solution and returned to the incubator for 90 min. After this resting period, short-term memory retention was assessed (trial 6) using the same procedure as the acquisition trials but without reward. Finally, all bees were cold-euthanised at -20°C.

### 2.7 Statistical analyses

All statistical analyses and data visualizations were performed in R version 4.5.1 (R Core Team, 2024) similar to our previous studies (e.g., Laußer and Kurze, 2025; Mandlinger and Kurze, 2025). The entire dataset, R markdown code and output, including all model details, validations, pairwise comparisons, are available via Zenodo (https://doi.org/10.5281/zenodo.20233307).

To analyse the sublethal effects of SIL and its synergistic potential with modern insecticides (ACE, SFX), we fitted generalized linear mixed models (GLMMs) with a binomial distribution (logit link function) using the *glmmTMB* package (Brooks et al., 2017). The GRS models used treatment doses (numeric) and sucrose concentration (factor) as fixed effects, while the PER models (learning) used treatment doses and trial numbers (factor). Because the negative controls (Table 1) were not significantly different for either the GRS (gustatory responsiveness) and PER learning assays (see results; Figs. 1a, 2a), they were pooled for all models including ACE and SFX. Interaction terms were included to specifically test for synergistic effects between SIL and insecticides on both gustatory responsiveness and PER learning over the course of each assay. All models included the bee ID, colony origin, and date as random effects to account for inter-individual (repeated measures), inter-colony, and temporal variability (including the different experimental blocks), respectively.

**Fig. 1.**
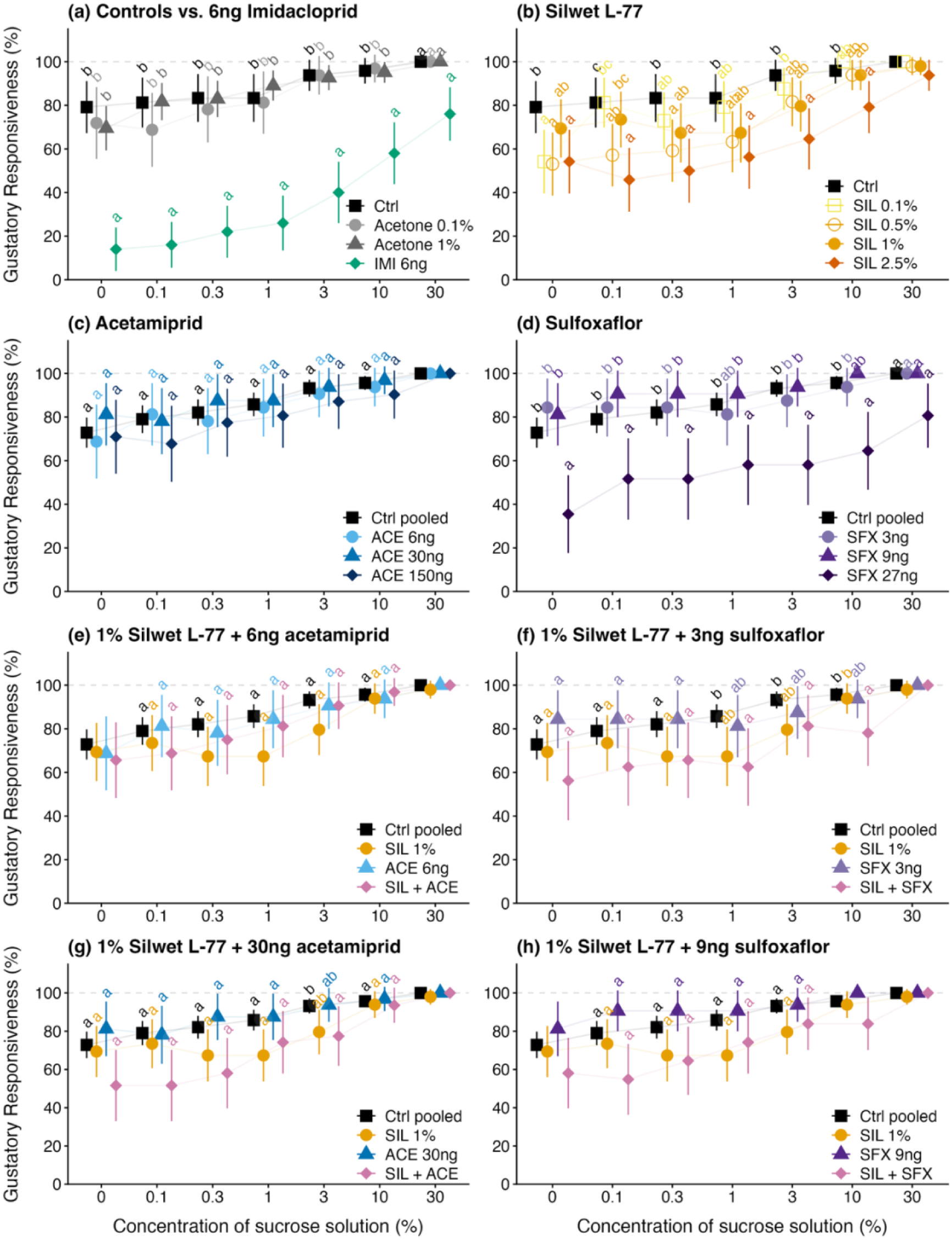
Effects of Silwet L-77 (SIL), acetamiprid (ACE), sulfoxaflor (SFX), and their interactions on the gustatory responsiveness (% mean GRS ± 95% CI) in honeybee foragers. *(a)* Validation of experimental procedure by comparing negative controls (Ctrl, acetone controls) and positive control (6 ng imidacloprid, IMI). Impact of *(b)* SIL, *(c)* ACE, *(d)* SFX alone. Interactions between 1% SIL and (*e, g*) ACE or (*f, h*) SFX. Results for interactions between 0.5% SIL and these pesticides are show in supplementary Fig. S2. Sample sizes n ≥ 31 bees per treatment (for details see Table 1). Treatment groups sharing a letter are not significantly different (*p* > 0.05). Ctrl = Control.

**Fig. 2.**
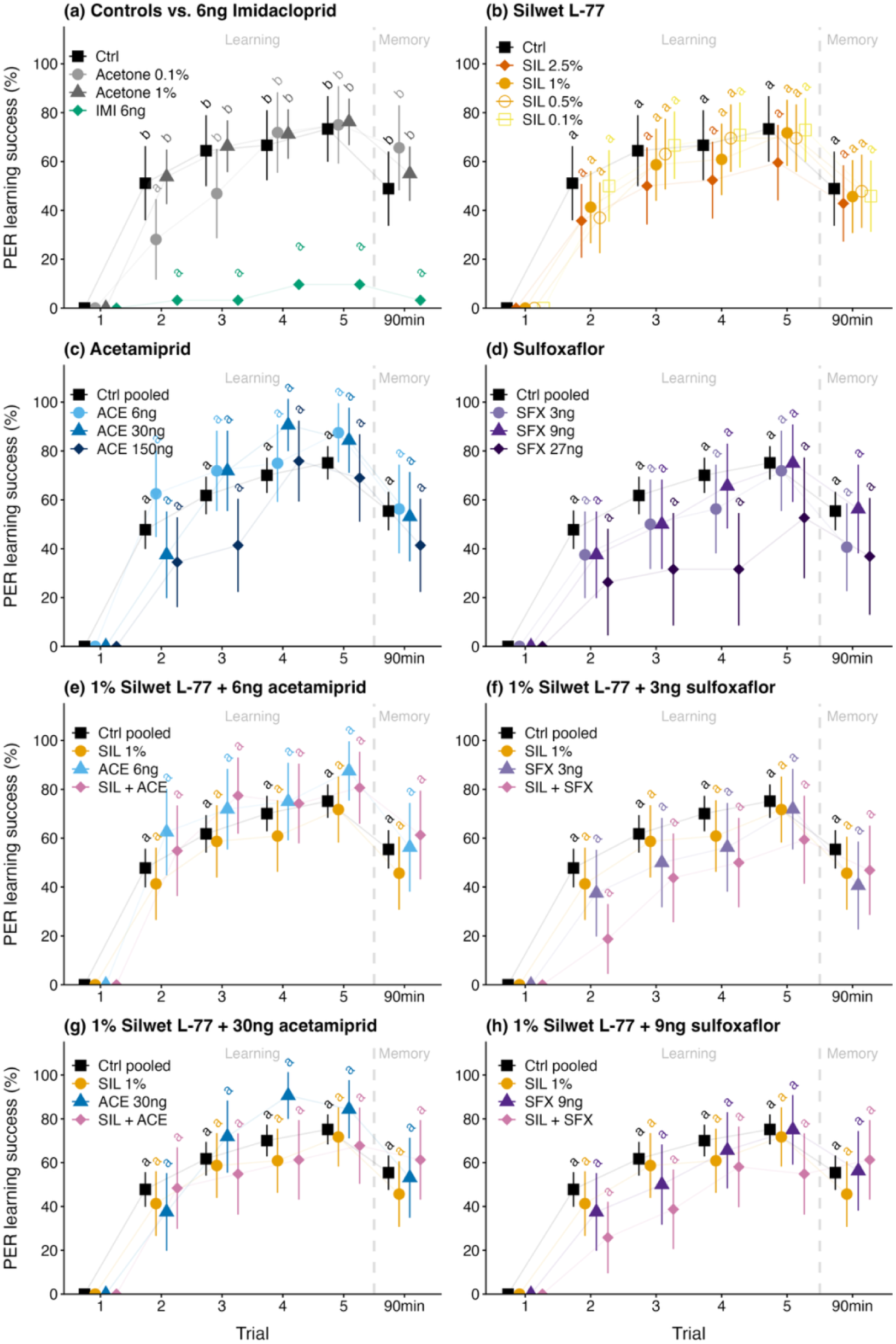
Effects of Silwet L-77 (SIL), acetamiprid (ACE), sulfoxaflor (SFX), and their interactions on associative learning (trials 1-5) and short-term memory after 90 min (% mean PER learning success ± 95% CI) in honeybee foragers. *(a)* Validation of experimental procedure by comparing negative controls (Ctrl, acetone controls) and positive control (6 ng imidacloprid, IMI). Impact of *(b)* SIL, *(c)* ACE, *(d)* SFX alone. Interactions between 1% SIL and (*e, g*) ACE or (*f, h*) SFX. Sample sizes n ≥ 31 bees per treatment (all except 27 ng SFX, n = 24; for details see Table 1). Treatment groups sharing a letter are not significantly different (*p* > 0.05). Ctrl = Control.

Model assumptions, including overdispersion and residual distribution, were validated using the *DHARMa* package (Hartig, 2020). To ensure model stability and convergence, specific sucrose concentrations (GRS) or trials (PER) had to be excluded from GLMM analysis when, for example, either all or no bees responded. These instances are indicated in the figures by the absence of pairwise comparisons, although their means and 95% confidence intervals (CI) are still displayed for descriptive purposes. Statistical significance (p < 0.05) was determined using the *Anova* function (Type II Wald χ² tests). Pairwise comparisons were conducted on treatment groups (factors) and odds ratios (OR) were calculated using *emmeans* (Lenth and Lenth, 2018) with Holm-Bonferroni corrections for multiple testing.

To test for overall treatment effects on mortality and sucrose unresponsiveness (failure to respond to 30% sucrose prior the olfactory PER conditioning), Fisher’s Exact Tests with 10,000 Monte Carlo simulations were performed. If significant, post-hoc pairwise Fisher’s Exact Tests were performed comparing selected treatment groups to the controls with Holm-Bonferroni corrections for multiple testing.

## 3. Results

### 3.1. Mortality

Mortality was generally low with 15 out of 855 (1.8%) individual bees dying during the entire experiment (Table 1). Nonetheless, overall mortality differed significantly across treatment groups (Fisher’s Exact Test with 10,000 Monte Carlo simulations, *p* < 0.0001). Strikingly, 25.0% (8 of 32) individuals treated with 27 ng SFX died in course of the experiment, which was significantly higher compared to the pooled controls, where only 0.6% (1 of 162) individuals died (Pairwise Fisher’s Exact Test with Holm-Bonferroni correction, *p* < 0.0001, odds ratio (OR) = 51.91, 95% CI: 6.49-2360.33). The second highest mortality occurred in the ACE 150 ng treatment with 6.25% (2 of 32) of individuals dying. However, this was not significantly different from the pooled controls (*p* = 0.071, OR = 10.52, 95% CI: 0.53-634.10).

### 3.2 Gustatory Responsiveness (GRS)

Analysis of the control data, including the positive control (6 ng IMI) in contrast to the negative controls (Ctrl, Acetone Ctrls), indicated a highly significant treatment effect on the GRS in honeybees (GLMM: χ² = 36.969, df = 3, *p* < 0.001; Fig. 1 a). Furthermore, the probability of unconditioned PER increased with increasing sucrose concentrations (χ² = 62.718, df = 6, *p* < 0.0001). Pairwise comparisons revealed that IMI significantly reduced gustatory responsiveness across sucrose concentrations from 0-10% (all *p* < 0.0001). No significant differences were detected among the negative control treatments (all *p* ≥ 0.555). All pairwise comparisons are available via Zendo (https://doi.org/10.5281/zenodo.20233307).

There was a significant effect of SIL concentration (χ² = 12.182, df = 1, *p* < 0.001; Fig. 1 b), sucrose concentration (χ² = 85.342, df = 5, *p* < 0.0001), and their interaction on the bee’s response (χ² = 16.817, df = 5, *p* = 0.005). While there was no significant difference in the bee’s response when presented with 30% sucrose across all SIL concentrations (all *p* = 1.0), pairwise comparisons revealed that the highest concentration (2.5% SIL) significantly reduced GRS relative to the control at most sucrose concentrations (0%: *p* = 0.010; 0.1%: *p* < 0.001; 0.3%: *p* < 0.001; 1%: *p* = 0.004; 3%: *p* < 0.001; 10%: *p* = 0.014). While 0.5% SIL differed from the controls at low sucrose concentrations (0%: *p* = 0.014; 0.1%: *p* = 0.021, and 0.3%: *p* = 0.020; 1%: *p* = 0.064; 3%: *p* = 0.118; 10%: *p* = 1.000), there was no significant difference between 1% SIL and the control (0-10%: all *p* ≥ 0.126). Furthermore, GRS did not differ significantly between the 0.5% and 1% SIL groups (0-10%: all *p* ≥ 0. 342) and did not differ consistently between the 1% and 2.5% groups (all *p* ≥ 0.118, except for 0.1%: *p* = 0.022 and 10 %: *p* = 0.065). Comparing the odds ratios (OR) indicated a dose-dependent decrease in likelihood of responding, for instance, to 0.3% sucrose with increasing SIL concentrations (0.1% SIL: OR = 10; 0.5% SIL: OR = 46; 2.5 % SIL: OR = 171), a pattern that was consistent across concentrations from 0.1-10% sucrose.

No significant treatment effect of acetamiprid on GRS was found (χ² = 0.547, df = 3, *p* = 0.908; Fig. 1 c). In contrast, SFX dose significantly affected GRS (χ² = 9.904, df = 1, *p* = 0.002; Fig. 1 d), but without an interaction effect between SFX dose and sucrose concentration (χ² = 7.372, df = 1, *p* = 0.194). While there was again no significant difference in response at 30% sucrose across all SFX doses (all *p* = 1.0), pairwise comparisons indicated a clear dose-response relationship at lower thresholds. For example, at 0.3% sucrose, control bees showed a drastically higher likelihood of responding than those treated with the highest SFX dose (27 ng SFX: OR = 1120, *p* = 0.002). In contrast, the lower SFX doses did not differ from the control at 0.3% sucrose (3 ng SFX: OR = 1, *p* = 0.991; 9 ng SFX: OR < 0.01, *p* = 0.760). Specifically, the highest SFX dose (27 ng) resulted in a significantly lower GRS than the controls (0%: *p* < 0.001; 0.1%: *p* = 0.005; 0.3%: *p* = 0.002; 1%: *p* = 0.004; 3% and 10%: *p* < 0.0001) and individuals treated with 3 ng (0%: *p* < 0.001; 0.1%: *p* = 0.007, 0.3%: *p* = 0.007; 1%: *p* = 0.070; 3%: *p* = 0.015; 10%: *p* = 0.013) or 9 ng SFX (0%: *p* < 0.001; 0.1%: *p* = 0.005; 0.3%: *p* = 0.004; 1%: *p* = 0.016; 3%: *p* = 0.005; but 10%: *p* = 1.0). No significant differences were observed between the 3 ng and 9 ng groups or between either group and the controls (0-10%: all *p* ≥ 0.360).

While we found no significant overall interaction effects on GRS between 0.5 % SIL and both insecticides (6ng ACE: χ² = 0.036, df = 1, *p* = 0.849, Fig. 1e; 30 ng ACE: χ² = 1.778, df = 1, *p* = 0.182, Fig. 1 g; 3 ng SFX: χ² = 1.068, df = 1, *p* = 0.301, Fig. 1f; 9 ng SFX: χ² = 0.113, df = 1, *p* = 0.737, Fig. 1h), there was a significant three-way-interaction effect with changing sucrose concentration for 6ng ACE (χ² = 11.825, df = 4, *p* = 0.019) and for 3 ng SFX (χ² = 11.923, df = 5, *p* = 0.036). There was also no significant interaction between 0.5% SIL and both insecticides (6 ng ACE: χ² = 1.756, df = 1, *p* = 0.185, Fig. S3a; 30 ng ACE: χ² = 0.445, df = 1, *p* = 0.505, Fig. S3c; 3 ng SFX: χ² = 0.283, df = 1, *p* = 0.594, Fig. S3b; 9 ng SFX: χ² = 1.018, df = 1, *p* = 0.313, Fig. S3d). Descriptive ORs revealed higher responsiveness of the controls in comparison to 1% SIL-pesticide mixtures, often exceeding the baseline effects of the insecticides alone (but mostly not SIL alone) such as at the critical 0.3% sucrose threshold (combinations of 1% SIL with 6 ng ACE: OR = 1.28; 30 ng ACE: OR = 59.172; 3 ng SFX: OR = 32.457; 9 ng SFX: OR = 2.111). A similar pattern was found for adjuvant-pesticide mixtures containing 0.5% SIL and either ACE or SFX (6 ng ACE: OR = 2.188; 3 ng SFX: OR = 16.737; with 9 ng SFX: OR = 6.000), but this was not the case for mixtures containing 0.5% SIL and 30 ng ACE (OR = 0.333). Importantly, these pairwise deviations from the controls predominantly reflect the independent gustatory suppression of SIL alone rather than an interactive, synergistic effect.

### 3.3 Sucrose responsiveness and exclusion rates

Across the entire experiment, only 25 out of 849 (2.9%) individual bees were unresponsive to the 30% sucrose stimulus. However, the proportion of unresponsive bees differed significantly across treatment groups (Fisher’s Exact Test with 10,000 Monte Carlo simulations, *p* < 0.0001). Most notably, 24.0% (12 of 50) of individuals treated with 6 ng IMI (positive controls) were unresponsive, which was significantly higher compared to the pooled negative controls (0 of 162, i.e. all responded) (Pairwise Fisher’s Exact Test with Holm-Bonferroni correction, *p* < 0.0001). Similarly, 19.4% (6 of 31) of bees treated with 27 ng SFX responded significantly less common to the 30% sucrose stimulus (*p* < 0.0001). Lastly, 6.25% (3 of 48) of individuals treated with 2.5% SIL were unresponsive to 30% sucrose (*p* = 0.012).

### 3.4 PER assay for associative learning and short-term memory

Analysis of the control data confirmed successful conditioning to the hexanal odour (GLMM: χ² = 37.092, df = 3, *p* < 0.0001; Fig. 2 a). Furthermore, treatment significantly affected both associative learning (χ² = 45.281, df = 3, *p* < 0.0001) and short-term memory (χ² = 13.638, df = 3, *p* = 0.003). Pairwise comparisons revealed that IMI (6 ng) significantly impaired associative learning and short-term memory compared to the negative controls across trials (e.g. for Ctrl: trial 2: *p* = 0.001; trial 3: *p* = 0.0001; trial 4: *p* < 0.0001; trial 5: *p* < 0.001; memory: *p* = 0.007). No significant differences were detected among negative control treatments (all *p* ≥ 0.109), except in a single time point (trial 2), where bees fed 0.1% acetone showed reduced learning compared to those fed pure sucrose solution (*p* = 0.042) or 1% acetone (*p* = 0.022) and did not differ significantly from the IMI group (*p* = 0.078). This validates the sensitivity of the PER assay to known neurotoxic stressors and consistency of the negative controls.

Overall SIL had a significant effect on associative learning (χ² = 5.849, df = 1, *p* = 0.016, Fig. 2 b), but without any significant differences between treatment groups across acquisition trials (all *p* ≥ 0.561). Short-term memory remained unaffected by SIL alone (χ² = 0.500, df = 1, *p* = 0.479). Evaluating the odds ratios (OR) indicated a potential dose-dependent effect, where controls showed a higher likelihood of learning and short-term memory retention compared to increasing SIL concentrations (e.g. at trial 3: 0.1% SIL: OR = 0.808; 0.5% SIL: OR = 1.083; 1% SIL: OR = 1.388; 2.5 % SIL: OR = 3.227), a pattern that was consistent across the acquisition and retention trials. Similarly, ACE and SFX alone also significantly affected learning overall (ACE: χ² = 4.991, df = 1, *p* = 0.025, Fig. 2 c; SFX: χ² = 5.305, df = 1, *p* = 0.021, Fig. 2 d), but not short-term memory (ACE: χ² = 2.611, df = 3, *p* = 0.456; SFX: χ² = 1.414, df = 1, *p* = 0.234). Pairwise comparisons revealed no significant effects of APC and SFX on learning (APC: all *p* ≥ 0.080; SFX: all *p* ≥ 0.057). However, descriptive OR comparison indicated a dose-dependent effect on learning for both APC (e.g. at trial 3: pooled Ctrl vs. 6 ng: OR = 0.594, 30 ng: OR = 0.676, 150 ng: OR = 5.26) and SFX (e.g. at trial 3: pooled Ctrl vs. 3 ng: OR = 1.469, 9 ng: OR = 1.349, 27 ng: OR = 5.800).

Finally, no significant interactions on learning and short-term memory were found between 1% SIL and ACE (learning (6 ng): χ² = 0.000, df = 1, *p* = 0.992; (30 ng): χ² = 0.720, df = 1, *p* = 0.396; memory (6 ng): χ² = 0.655, df = 1, *p* = 0.419; (30 ng): χ² = 1.158, df = 1, *p* = 0.282, Fig. 2 e, g), nor between 1% SIL and SFX (learning (3 ng): χ² = 0.886, df = 1, *p* = 0.347; (9 ng): χ² = 1.047, df = 1, *p* = 0.306; memory (3 ng): χ² = 0.884, df = 1, *p* = 0.347; (9 ng): χ² =1.047, df = 1, *p* = 0.306; Fig. 2 f, h). This lack of significance was also consistent for interactions between 0.5% SIL and ACE (learning (6 ng): χ² = 2.233, df = 1, *p* = 0.135; (30 ng): χ² = 0.578, df = 1, *p* = 0.447; memory (6 ng): χ² = 0.048, df = 1, *p* = 0.827; (30 ng): χ² = 1.202, df = 1, *p* = 0.273, Fig. S3 a, c), and for 0.5% SIL and SFX (learning (3 ng): χ² = 2.352, df = 4, *p* = 0.671; (9 ng): χ² = 1.991, df = 4, *p* = 0.737; memory (3 ng): χ² = 2.076, df = 1, *p* = 0.651; (9 ng): χ² = 1.991, df = 4, *p* = 0.737; Fig. S3 b, d). Descriptive OR comparisons against the controls at for instance trial 3, further reflect the absence of distinct interactive effects between and SIL and ACE (6 ng ACE + 1% SIL: OR = 0.461; 30 ng ACE + 1 % SIL: OR = 2.841; 6 ng ACE + 0.5% SIL: OR = 2.186; 30 ng ACE + 0.5% SIL: OR = 0.767). However, for the combination of SIL and SFX, OR comparisons revealed a visible trend toward lower learning success, despite the overall model interaction being statistically non-significant (3 ng SFX + 1% SIL: OR = 2.478; 9 ng SFX + 1% SIL: OR = 3.179; 3 ng SFX + 0.5% SIL: OR = 3.266; 9 ng SFX + 0.5% SIL: OR = 2.476).

## 4. Discussion

Our study provides early evidence for the sublethal effects of oral exposure to both single compounds (SIL, ACE, and SFX) and adjuvant-insecticide mixtures (SIL-ACE and SIL-SFX) on gustatory responsiveness, PER learning performance, and short-term memory in honeybee foragers under standardised laboratory conditions. While organosilicone adjuvants like SIL are widely used to enhance spraying efficiency and pose direct risk by increasing cuticle penetration following spray contact (Mullin et al., 2016; Shannon et al., 2023; Straw et al., 2022), foragers may also ingest these mixtures via contaminated nectar and pollen. However, the sublethal effects of ingested adjuvants and their interaction with insecticides on bee physiology and the intestinal epithelium remain poorly understood. While we found that high-dose single-compound exposure to 2.5% SIL (Fig. 1 b) and 27 ng SFX (Fig. 1 d) significantly reduced gustatory responsiveness (GRS), there were no significant interaction effects for SIL-ACE nor SIL-SFX mixtures at these potentially field-relevant doses (0.5% or 1% SIL combined with 6 or 30 ng ACE, or 3 or 9 ng SFX) (Fig. 1 e-h; Fig. S2 a-d). Similarly, while single-compound exposure to SIL, ACE, and SFX significantly affected associative PER learning without impacting short-term memory retention after 90 minutes (Fig. 2 b-d), the tested adjuvant-insecticide mixtures showed no significant interaction effects on learning performance and short-term memory (Fig. 2 e-h; Fig. S3 a-d).

Overall, our results from negative and positive controls (Figs. 1 a, 2a) aligned with previous studies (e.g., Ciarlo et al., 2012; Scheiner et al., 2013), validating the sensitivity and reliability of the methods used. Although mortality was generally low (1.8%) and comparable across treatment groups (Table 1), bees exposed to 27 ng SFX died disproportionately more often (25%) during the experimental procedure. This mortality rate is consistent with previously reported oral toxicity levels for SFX, where 30 ng/bee induced similar mortality after 48 h and the LD_50_ was estimated at 95.88 ng/bee (Cartereau et al., 2022). Another study reported an LD_50_ of 145.43 ng/bee at 3 h and 55.38 ng/bee at 24 h post-exposure (Azpiazu et al., 2019).

Although co-formulants in tank-mixtures are often labelled as ‘inert’, our data show that SIL significantly reduces GRS, particularly at 2.5% (Fig. 1 b). While we also found that SIL significantly affected associative PER learning overall, pairwise comparisons revealed no significant differences between treatment groups. This result contrasts with findings by Ciarlo et al. (2012), who showed that consuming 1% SIL, among other organosilicone adjuvants (Dyne-Amic^®^, Syl-Tac^®^, and Sylgard^TM^ 309), significantly impaired olfactory learning in honeybees. A possible explanation for this discrepancy might be differences in exposure volume. In their study, bees fed freely from a saturated cotton swab, potentially resulting in a higher total ingested dosage. Their longer starvation period of 18 h may have also increased physiological stress. This idea is supported by a recent study showing that starvation combined with pesticide exposure can overwhelm antioxidant defences in honeybees (Hernández-Rivera et al., 2025).

While sublethal doses of ACE did not significantly affect GRS (Fig. 1 c), SFX at 27 ng/bee significantly reduced sucrose responsiveness (Fig. 1 d). Overall, both insecticides significantly impaired associative learning across trials (Fig. 2 c, d). Although descriptive ORs indicated dose-dependent reductions in learning performance, post-hoc pairwise comparisons revealed no significant differences between treatment groups. In part, this discrepancy likely reflects a combination of reduced statistical power following conservative Holm-Bonferroni adjustments and the exclusion of unresponsive individuals prior to PER conditioning. Indeed, sample sizes for the 27 ng SFX treatment group were significantly reduced by 31.25% (to only 22 of 32 bees; Table 1) due to increased mortality and the exclusion rule, introducing survivorship bias. Regardless, these findings are largely consistent with the literature reporting potential risks of ACE and SFX to honeybees and other insect pollinators (e.g., DesJardins et al., 2023; Mandal et al., 2026; Spence et al., 2026). For example, feeding 15 ng SFX to honeybees impaired learning and altered the expression of nAChR subunits in the brain (Cartereau et al., 2022). Similarly, SFX was shown to drastically reduce homing ability at 26 ng/bee, whereas ACE (up to 61 ng/bee) had no significant effect (Capela et al., 2022). In our study, only the highest exposure scenario (150 ng/bee) indicated a negative trend in learning performance, characterised by a decreased likelihood of successful learning, whereas bees exposed to 6 ng and 30 ng ACE remained unaffected. Earlier studies reported that acute ACE exposure altered gustatory responsiveness (El Hassani et al., 2008), while chronic exposure for 11 days (100 and 1000 ng/bee/day) increased responsiveness to water without impairing learning performance (Aliouane et al., 2009). Higher doses of ACE (100 ng/bee) caused a delayed impairment of short-term memory retention without effects on learning (Thany et al., 2015). Finally, chronic exposure to field-relevant concentrations of ACE (12 ng and 120 ng/bee/day fed for one week) showed no significant effect on sucrose responsiveness or olfactory learning (Schuhmann and Scheiner, 2023).

Contrary to our initial hypothesis, we found no significant synergistic interactions between the adjuvant SIL and either ACE or SFX regarding sensory perception (Fig. 1 e-h, Fig. S2 a-d), or olfactory learning and short-term memory performance for the tested dose combinations (Fig. 2 e-h, Fig. S3 a-d). At first glance, this is surprising, given that adjuvants like SIL have been shown to increase the acute toxicity of insecticides like ACE. For example, the LD_50_ of spray exposure to ACE alone was 26 µg/bee, but in combination with SIL, the LD_50_ dropped to < 4.79 µg/bee (Chen et al., 2019). Similarly, ACE applied in spray formulations with the crop oil adjuvant LI 700 and the organosilicone Break-Thru^®^ S 301 significantly increased mortality in honeybees (Wernecke et al., 2022). These findings suggest that single oral exposure events of adjuvant-insecticide mixtures, as used in our study, may not exhibit the same synergistic potency as lethal sprayed doses. Thus, SIL might either not amplify the impact of ACE and SFX on neurobiological pathways at our chosen concentrations, or the lethal synergistic effects observed in spray experiments may mainly be driven by enhanced spreading of insecticides across the bee’s cuticle and potentially increased spiracular penetration (Shannon et al., 2023). It is also plausible that the interaction effects observed in mortality assays stem from a reduction in respiratory capacity, as these agrochemicals may block the tracheal system, leading to suffocation (Straw et al., 2022).

We speculate that higher dose combinations or chronic/repeated exposure to SIL-SFX mixtures would result in significantly reduced gustatory responsiveness and learning performance, similar to the higher doses of 2.5% SIL and 27 ng SFX alone (Fig. 1 b, d; Fig. 2 b, d). Indeed, descriptive OR comparisons at trial 3 already indicate a downward trend in learning performance for the SIL-SFX mixtures, where bees consistently showed a decreased likelihood of learning success relative to controls (e.g., 30 ng ACE + 1 % SIL: OR = 2.841; 9 ng SFX + 1% SIL: OR = 3.179), although this was not statistically significant. Despite a potential limitation in statistical power, this descriptive decline points to an additive negative effect rather than a strong synergistic interaction. Similarly, while there were no significant interaction effects of the adjuvant-insecticide mixtures on the sucrose responsiveness in foragers, descriptive ORs indicated gustatory suppression at the 0.3% sucrose threshold compared to the controls (e.g., 30 ng ACE + 1% SIL: OR = 59.172; 3 ng SFX + 1% SIL: OR = 32.457).

Regardless, the reduction in sucrose responsiveness caused by potential field-relevant doses of SFX and SIL alone may have important consequences for honeybees, serving as an initial risk indicator for other insect pollinators. Gustatory responsiveness is closely linked to foraging motivation, reward evaluation, and division of labour in honeybees (Scheiner, 2004; Scheiner et al., 2013). This emphasises the importance of including GRS along with PER conditioning and memory retention assays in pesticide risk assessment. Standard regulatory assessments are still largely based on single compounds and acute mortality measurements in honeybees, which likely underestimate the sublethal, synergistic, and chronic effects, as well as widely unexplored effects in non-*Apis* insect species (Desneux et al., 2007; Straw et al., 2022; Tosi et al., 2022; Wintermantel et al., 2026). Our experiments focused on acute behavioural responses and therefore, do not account for delayed effects or chronic exposure. While laboratory behavioural assays do not reflect the complex cognitive challenges of the real world, they offer great value for the initial identification of sublethal effects before moving on to more complex tests, such as assessing bees’ foraging behaviour or navigation ability in the field (DesJardins et al., 2023).

## Conclusion

Although we found no evidence of synergistic effects of adjuvant-insecticide mixtures by acute oral exposure at the sensory-cognitive level, this does not imply that spray adjuvants are ‘inert’. Growing evidence suggests that organosilicone adjuvants alone can impair sensory perception and learning behaviour. Consequently, co-formulants in tank-mixtures should be considered in regulatory frameworks. Beyond routine acute mortality and risk assessments, incorporating sensory-cognitive assays (such as GRS and PER) will provide more comprehensive understanding of how sublethal doses impact on physiology and health of insect pollinators.

## Data accessibility statement

Dataset and all R scripts used for statistical analyses and producing all Figures are available from a Zenodo repository: https://doi.org/10.5281/zenodo.20233307. Electronic supplementary materials are available online.

## Statement of authorship

M.A. and N.B.: investigation, data curation. M.M.M. and J.O.: conceptualization, methodology, writing – review and editing; C.K.: conceptualization, methodology, investigation, data curation, resources, validation, formal analysis, writing – original draft preparation, writing – review and editing, visualization, supervision, project administration.

## Acknowledgments

We are grateful to Prof. Dr. Joachim Ruther for supplying sulfoxaflor and lending us the Syntech stimulus controller. We also thank three anonymous reviewers for helping us to improve the overall quality of our manuscript.

## Ethics statement

This study was conducted in accordance with the ethical regulations of the German Animal Welfare Act (TierSchG) for conducting experiments with insects.

## AI declaration

We used used Gemini (Google, 2026) for proofreading our manuscript and made adjustments when needed. We take full responsibility for the content of the published article.

