## Supplementary Figures for "Sublethal effects of the organosilicone surfactant Silwet L-77 and its interactions with acetamiprid and sulfoxaflor on honeybee (*Apis mellifera*) gustatory responsiveness and olfactory learning"

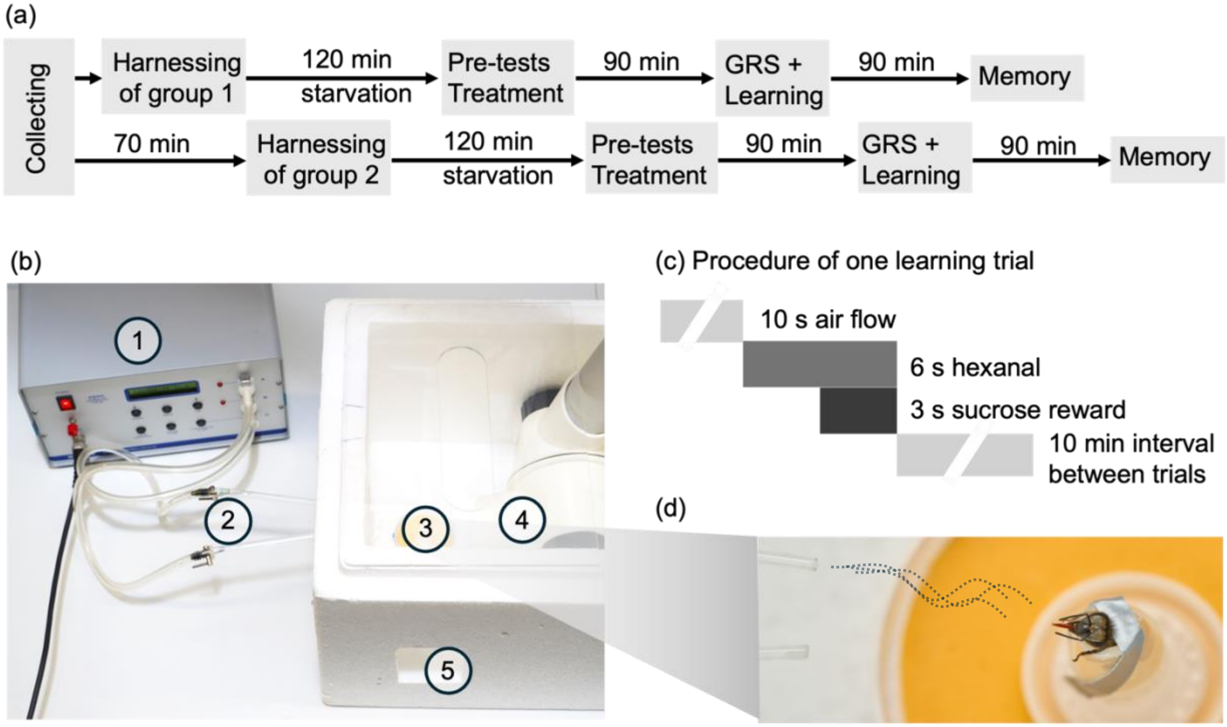


**Fig. S1. Overview of the experimental workflow and setup.** (a) Flow chart showing the experimental timeline for honeybee foragers collected from one of four hives each day. Bees were randomly assigned to one of two groups prior to double-blind treatment allocation. (b) The experimental setup consisted of the following components: (1) a stimulus controller (CS-55, Ockenfels Syntech GmbH) delivering a continuous stream of air (1.2 L $min^{-1}$) passing through (2) one of two Pasteur pipettes (one containing a 4 x 1 mm piece of filter paper with 4 µL hexanal), (3) the test bee placed about 4.5 cm in front of the of these pipettes (see panel d) inside an open polystyrene box, (4) a table fume hood (Siemtech Elektrotechnik Heuschmann GmbH) positioned behind the bee to exhaust the air stream, and (5) a small window (5.5 x 7 cm) through which the sucrose reward was provided. (c) Procedure for a single associative learning trial using the proboscis extension response (PER) assay. Between individual bees, the experimental chamber was vented for 18 s. The procedure was repeated four times per bee at 10-min intervals (totaling five conditioning trials). (d) Positive PER of a conditioned honeybee inside the experimental chamber during the first 3 s of hexanal exposure (testing phase; odour is indicated by dotted lines) without sucrose reward.

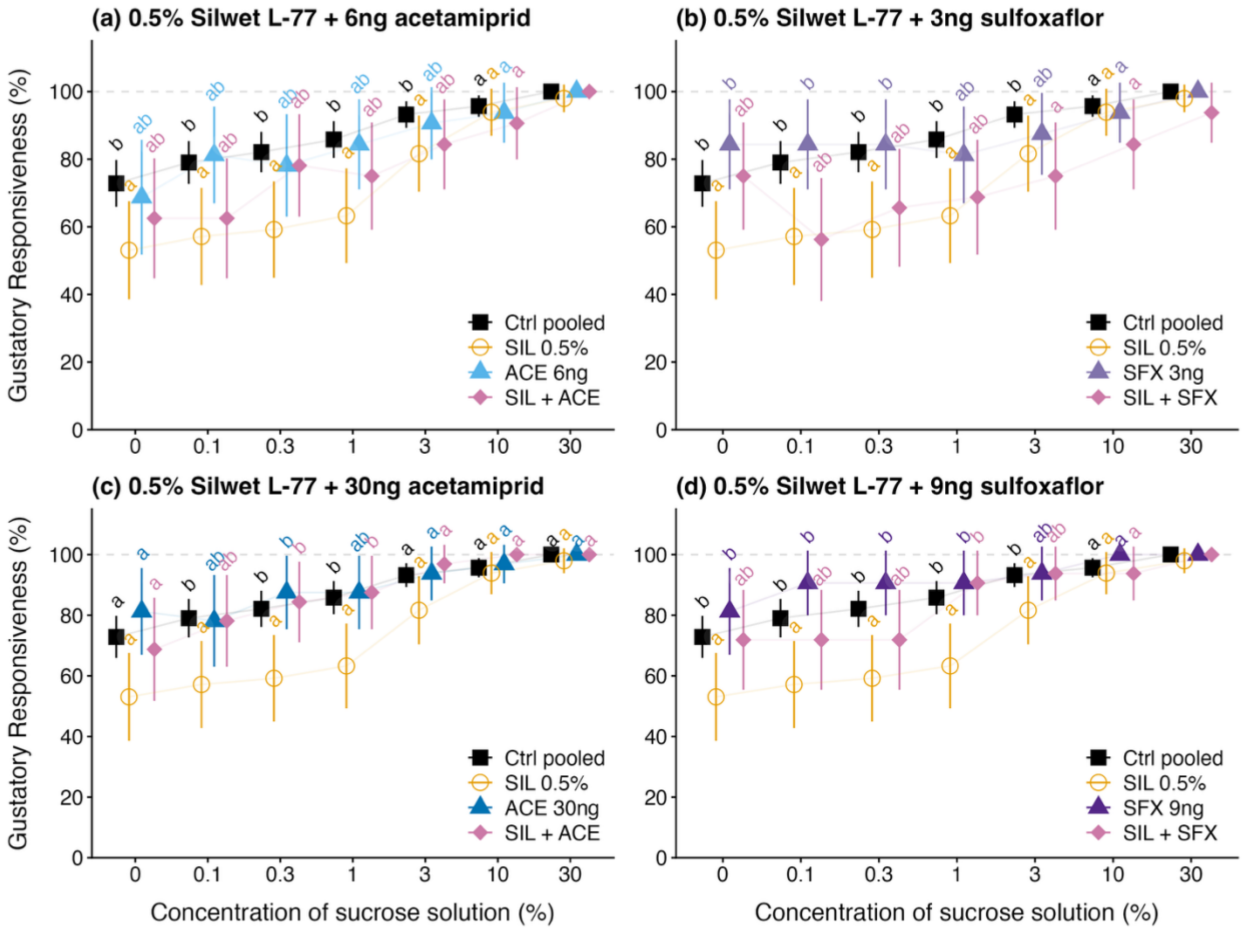


**Fig. S2. Synergistic effects of 0.5% Silwet L-77 (SIL) with acetamiprid (ACE; a, c) or sulfoxaflor (SFX; b, d) on the gustatory response score (mean GRS ± 95% CI) in honeybee foragers.** Sample sizes n ≥ 31 bees per treatment. Treatment groups sharing a letter are not significantly different (*p* > 0.05).

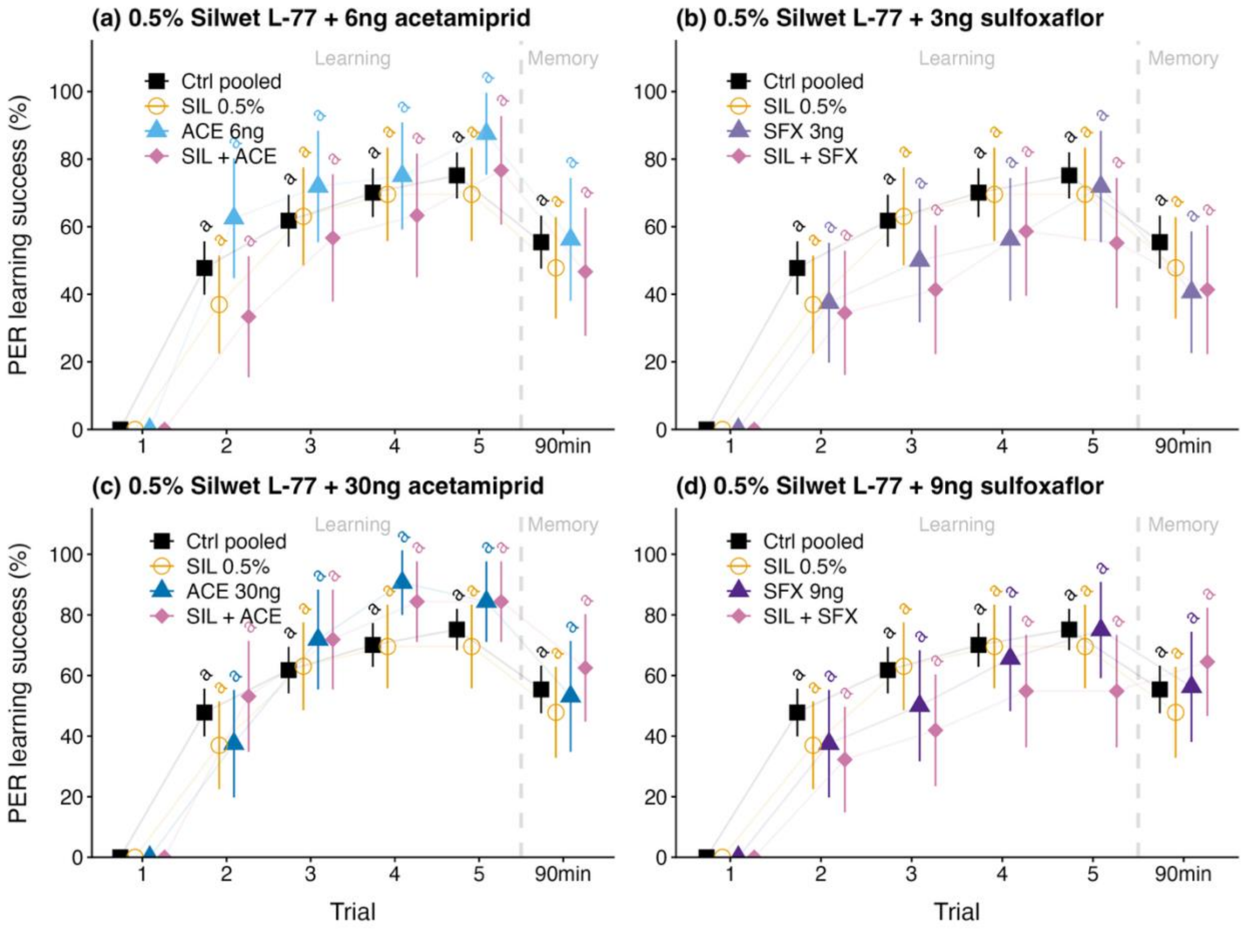


**Fig. S3. Synergistic effects of 0.5% Silwet L-77 (SIL) with acetamiprid (ACE; a, c) or sulfoxaflor (SFX; b, d) on associative learning (Trials 1-5) and short-term memory after 90 min (mean PER ± 95% CI) in honeybee foragers.** Sample sizes n ≥ 31 bees per treatment. Treatment groups sharing a letter are not significantly different (*p* > 0.05).
